# RNA-DCGen: Dual Constrained RNA Sequence Generation with LLM-Attack

**DOI:** 10.1101/2024.09.23.614570

**Authors:** Haz Sameen Shahgir, Md. Rownok Zahan Ratul, Md Toki Tahmid, Khondker Salman Sayeed, Atif Rahman

**Affiliations:** University of California, Riverside; Bangladesh University of Engineering and Technology

## Abstract

Designing RNA sequences with specific properties is critical for developing personalized medications and therapeutics. While recent diffusion and flow-matching-based generative models have made strides in conditional sequence design, they face two key limitations: specialization for fixed constraint types, such as tertiary structures, and lack of flexibility in imposing additional conditions beyond the primary property of interest. To address these challenges, we introduce RNA-DCGen, a generalized framework for RNA sequence generation that is adaptable to any structural or functional properties through straightforward finetuning with an RNA language model (RNA-LM). Additionally, RNA-DCGen can enforce conditions on the generated sequences by fixing specific conserved regions. On RNA generation conditioned on RNA distance maps, RNA-DCGen generates sequences with an average *R*^2^ score of 0.625 compared to random sequences that score only 0.118 over 250 generations as judged by a separate more capable RNA-LM. When conditioned on RNA secondary structures, RNA-DCGen achieves an average F1 score of 0.4 against a random baseline of 0.006.

## 1 Introduction

RNA plays a dual role in biological processes, firstly by acting as an information carrier from DNA to the ribosome in the form of mRNA, and secondly through functional roles such as RNA interference, riboswitches, and ribozymes. While the design of protein sequences with structural constraints has been significantly advanced due to the abundant availability of 3D structural data from the Protein Data Bank (PDB), the design of RNA sequences remains challenging due to RNA’s inherent structural flexibility, functional diversity, and the limited availability of high-resolution RNA data. The interest of generating RNA sequences can be elucidated by RNA-based therapeutics [16] [6], vaccine design, target design for gene-editing [24], gene silencing by RNA-interference.

The first attempt for RNA sequence generation was approached heuristically, aligning the RNA fragment motifs utilizing only the secondary structures of RNA [10] [26]. Such heuristics often suffer from the diverse large space of possible RNA sequences. [9] [15] showed that heuristics based on restrictive motif regions don’t fully capture the conformational dynamics of RNA functionality. The main challenge of RNA design using deep learning has been due to scarcity of RNA structural/functional data at nucleotide level [19]. RNA is ineherently more conformationally dynamic and the flexibility is driven by base pairing interaction between the strands and base stacking among the rings of spatial nucleotides [22]. Moreover, RNA nucleotides (analogous to residues in protein) contain 13 atoms in contrast to 4, causing 7 torsion angles to persist in each nucleotide’s point cloud, adding higher complexities to the structural design of RNA.

Recent work includes gRNAde [13], a geometric deep learning-based modeling for the generation of RNA sequences conditioned on 3D structure and RNA-FrameFlow [2], a flow-matching-based generative model for RNA 3D backbone design. However, these methods are primarily focused on structural constraints and lack the flexibility to incorporate additional conditions during sequence generation.

Due to limited data for RNA downstream tasks (typically a few hundred to a few thousand labeled sequences), RNA language models largely follow the pretrain-then-finetune paradigm. Most RNA language models use the bidirectional encoder-only transformer architecture [8]. They are typically pre-trained on general-purpose contexts using sequences from both coding and non-coding regions provided by the RNAcentral database [5]. Popular RNA language models include RNA-FM [4], RNA-MSM [23], RiNALMo [17] and BiRNA-BERT [20]. Similar to BERT [8], RNA language models learn deep contextual representations of RNA sequences that can be utilized for downstream tasks such as distance map prediction, secondary structure prediction, torsion angle prediction, etc. with relatively little data.

Unlike transformer-decoder-based generative models such as GPT [3] and LLaMA [21] in linguistics or ESM [11] for protein generation, encoder-based RNA language models are limited to predicting RNA properties, and generating sequences from these models becomes challenging when additional constraints must be applied to the sequence. As such, our key insight is to pose RNA generations as a search problem and use the predictive capabilities of RNA language models to effectively reduce the otherwise exponential (4^*N*^) search space of finding an RNA sequence of length *N*. Furthermore, we can combine arbitrary constraints during the search process as long as the RNA language model can predict said constraints effectively.

We draw inspiration from the domain of automated LLM jail-breaking and adapt the Greedy Coordinate Gradient Search algorithm [28] to encoder-only RNA language models, effectively transforming a nucleotide-level property prediction model into a generative model. Our method, RNA-DCGen, introduces a generalized framework that can be applied to RNA language models capable of predicting various molecular features, enabling the generation process to be conditioned both on the sequence itself and on specified constraints. This dual conditioning allows our approach to impose multiple requirements simultaneously during RNA sequence generation, enhancing its adaptability and applicability to diverse RNA design tasks.

## 2 Methods

In this section, we first discuss the complete working pipeline of RNA-DCGen (section 2.1) Then we discuss about the conditional properties that we impose while generating RNA sequences (section 2.2). Next, we discuss the statistical constraints that we can the generated sequences themselves with conserved regions (section 2.3). Finally, we describe the Gradient-based Greedy Search algorithm 2.4.

### 2.1 Complete Working Approach of RNA-DGCen

The overall pipeline of RNA-DGCen is represented in Figure 1. Firstly, we finetune an RNA language model (BiRNA-BERT [20] in RNA-DCGen) for the two constraints, RNA distance-map, and RNA secondary structure. Secondly, as the input to the generator, we provide the desired properties and a random input sequence. In the case of fixed or conserved generation, certain portions of the input sequence are kept fixed. Thirdly, the finetuned LLM generates predictions for the input sequence, and we compute the loss between the predicted properties and the desired properties. This loss is backpropagated through the LLM to generate a gradient distribution at each position for all possible words in the vocabulary. Next, we utilize a modified Gradient Coordinate Search [28] as presented in Algorithm 1 to calculate a gradient-guided search to modify certain portions of the sequence. Finally, to validate the quality of the generated sequence, we use another finetuned RNA language model (RiNALMo [17]) as an independent discriminator and compare the desired property quality between the ground properties and those of the generated sequences. We intentionally use the 650M parameter state-of-the-art RiNALMo for accurate validation and the 117M parameter, yet still capable, BiRNA-BERT for fast sequence generation. The two models differ in not only parameter count but also architectural choices and training data, which ideally reduces the risk of generating RNA sequences that aren’t biologically meaningful and exploit neural network weaknesses [27].

**Figure 1:**
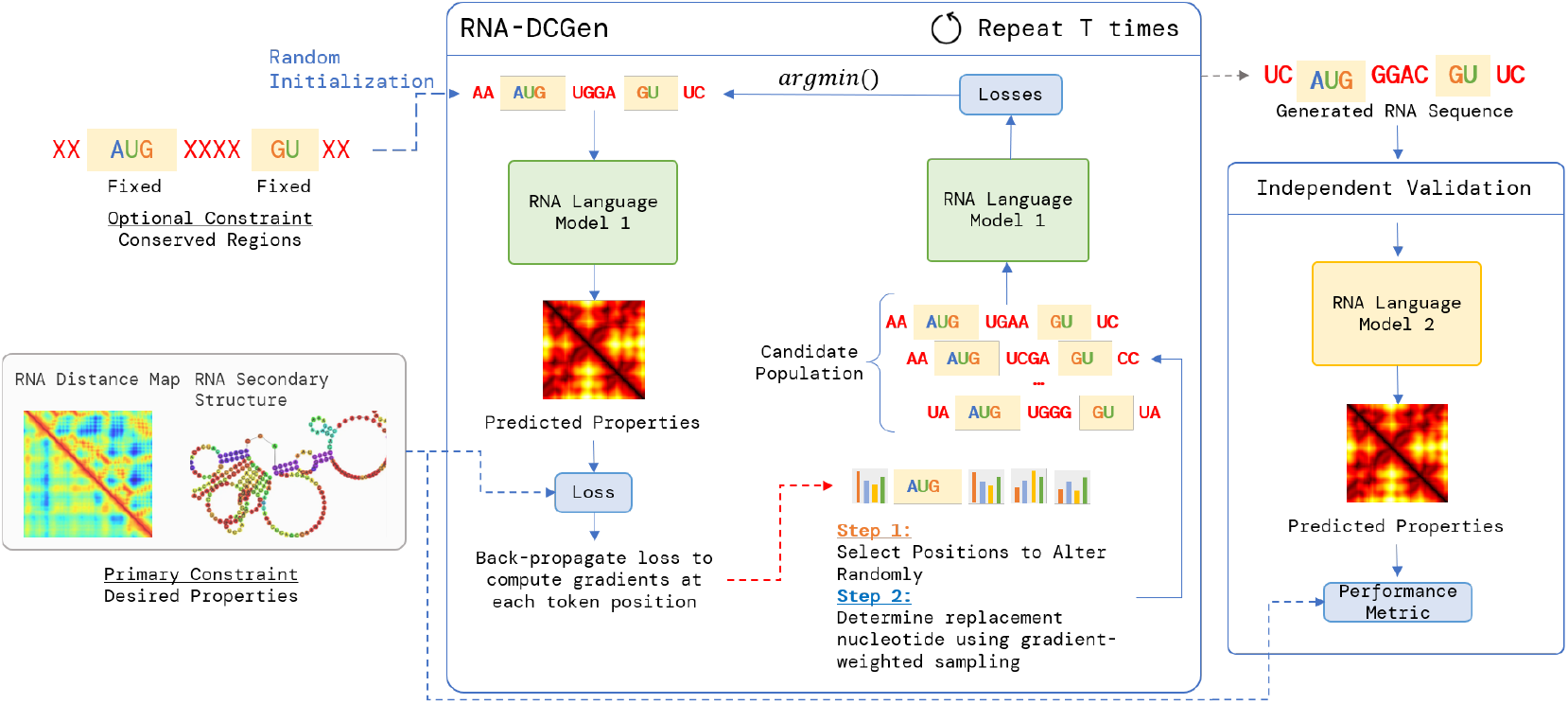
Overview of the proposed RNA-DCGen method. At each time step, the loss of the starting sequence is back-propagated to recover per-position, per-nucleotide gradients. Intuitively, a higher gradient for nucleotide *χ* at position *i* indicates that replacing the current nucleotide at *i* with *χ* will reduce the overall loss. This gradient information is then used to generate *B* new candidate RNA sequences from which the one with the lowest loss is kept for the following iteration. After *T* time steps, the final RNA sequence is generated. To ensure that the generated RNA sequence is biologically meaningful, a separate independent discriminator is used for validation.

### 2.2 Tasks and Datasets for Enforcing Conditional Properties

In this work, we experiment with two different types of property constraints: *RNA secondary structure* and *RNA distance map*. RNA-DCGen takes as input any of these conditional properties and generates sequences with these desired properties by backpropagating through the RNA-LM. The RNA distance map dataset is collected from trRosettaRNA’s webserver ^3^, and the secondary structure dataset is collected from the BEACON benchmark dataset of bpRNA ([18]).

### 2.3 Constraints on Generated Sequences: Full vs Conserved Regions

RNA-DCGen can generate the entire RNA sequence using property constraints or **optionally** fix arbitrary regions that will remain unchanged throughout the generation process. This allows the sequence designer to enforce the inclusion of biologically significant sub-sequences such as RNA aptamer binding sites [14, 12], crucial for molecular recognition, or guide RNA (gRNA) regions used in CRISPR-Cas systems [25] for precise genome editing. Fixing sequence regions helps ensure that docking sites or DNA targeting become more realizable while optimizing other parts of the sequence. In our experiments, we simulate biologically significant sub-sequences using the RNA tokenization statistics reported by BiRNA-BERT [20] and fix 3 of the most common sub-sequences per RNA which are at least 5 nucleotides long. We note that in real applications, the designer would manually design constraints based on specific applications of the generated RNA.

### 2.4 Sequence Generation from an LLM Adversarial Attack Optimization Perspective

The authors of LLM-Attack [28] showed that it’s possible to force LLMs to generate a response starting an affirmation (e.g. *“Sure, here is*…*”*) by appending an adversarial suffix to the end of an LLM query. The adversarial suffix is optimized by computing the gradient of the loss with respect to the desired affirmative response and using the gradient information to search for effective adversarial suffixes using the Gradient-Coordinate Search (GCG) algorithm. Our key insight is that this prompt optimization process is arbitrary; the entire sequence and not just the suffix can be optimized and any loss function can be used. We therefore adopt the GCG algorithm to generate prompts (RNA sequences in the case of RNA-DCGen) that match a desired property using RNA language models,

#### Algorithm 1: Gradient-based Greedy Search

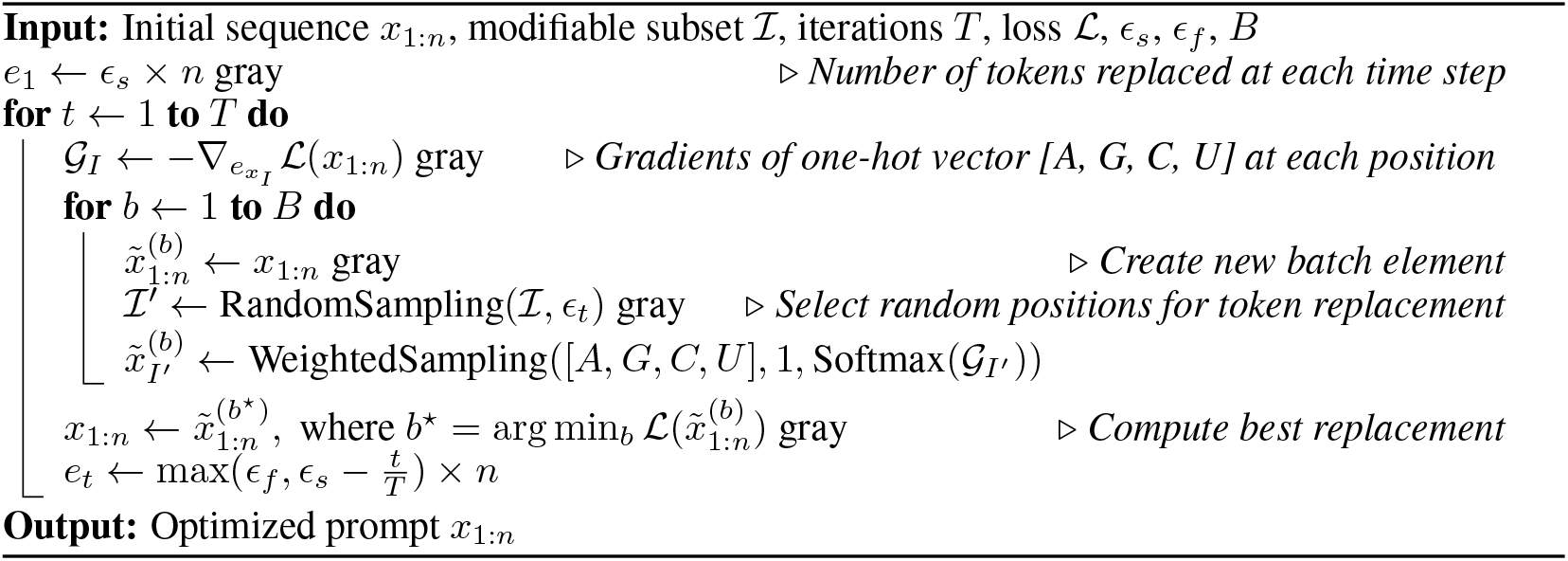

## 3 Results

In this section, we discuss the qualities of the generated sequences for both fixed generation and full generation. We consider two properties of RNAs while imposing constraints: RNA distance map and secondary structure.

### 3.1 Performance of RNA-DCGen with constraints on RNA Distance Map

The distance map provides a way to impose conditions on the pairwise distances for each nucleotide of the RNA sequence. In Table 1, we show the quality of the generated sequences with conditions on the distance map. For *statistically fixed generation*, the R2 score between the desired properties and the predictions from the discriminator for ground sequences (sequences with which the language models are trained) is 0.810, which is 0.693 for the sequences generated by RNA-DCGen.

**Table 1:** Comparison of the Performance of RNA-DCGen in sequence generation, conditioned on RNA Distance Map and Secondary Structure under different conditions.

|  | Condition on RNA Distance Map (R2 Score) |  | Condition on RNA Secondary Structure (F1 Score) |  |
| --- | --- | --- | --- | --- |
|  | Statistically Fixed | Full Generation | Statistically Fixed | Full Generation |
| Ground | 0.810 | 0.810 | 0.729 | 0.729 |
| Generated | 0.693 | 0.625 | 0.513 | 0.400 |
| Random | 0.257 | 0.118 | 0.005 | 0.006 |

We compare the performance with randomly generated sequences of similar length, and a R2 score of 0.257 is observed, which indicates that the generated sequences with RNA-DCGen indeed preserve the structural constraint imposed by the distance map. In Figure 2 (a), we demonstrate a representative sequence that is generated under the statistically fixed generation scheme, with an R2 score of 0.60. For *full sequence generation*, the average R2 score of the generated sequences is 0.625 indicating that it is challenging to generate the complete sequence under the constraints when no portion of the input sequence is kept fixed.

**Figure 2:**
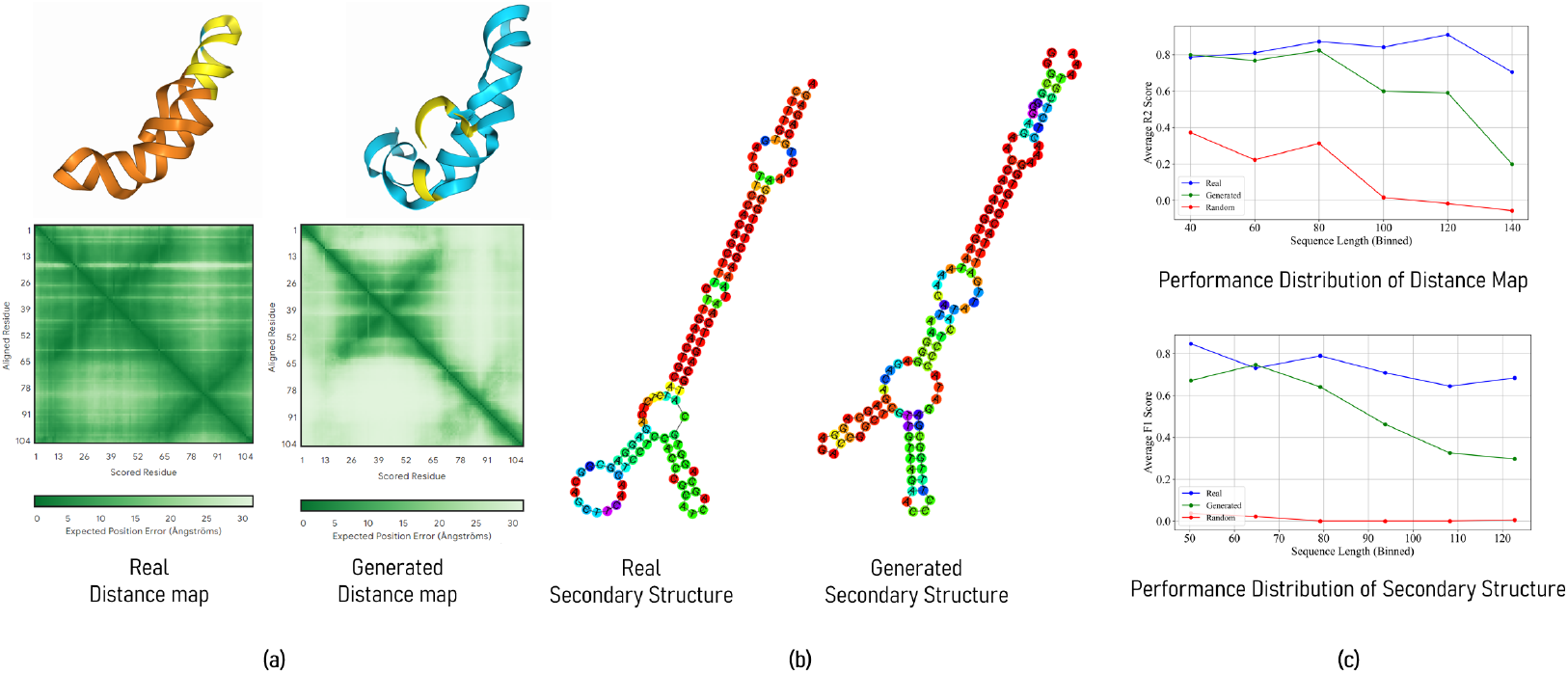
(a) Alphafold-3 [1] distance map predictions for ground-truth vs generated sequence with R2 score of 0.60 and sequence length of 104 (b) Real vs generated sequence’s secondary structure with F1 score of 0.79 and sequence length 126. (c) Influence of sequence length on the quality of the generated sequences. For shorter sequences, the property conditioned generation performance shows a strong correlation with the ground truth and the performance quality gradually deteriorates for much longer sequences.

### 3.2 Performance of RNA-DCGen with Constraints on RNA Secondary Structure

Secondary structure is a 2D property of RNA structures that determines which nucleotides within the sequences will form different types of loops. In Table 1 we see that, for ground sequences under the *statistically fixed generation*, the F1 score is 0.729 whereas for the generated sequences this score is 0.513. It is to be noted that, secondary structure prediction from a sequence-only model is itself a challenging task where the F1 score for forward prediction with RiNALMo [17] is 0.73. Thus, the generative task based on this trained language model also provides a lower F1 score. Random sequences generated under the desired secondary structure condition also achieve a score of 0.005 indicating that RNA-DCGen well captures the conditional signals on secondary structure. In Figure 2 (b), we demonstrate the outcome of a generated sequence under a conditioned secondary structure with an F1 score of 0.79 and a sequence length of 126. In Table 1, we observe a similar pattern to the distance map where *full sequence generation* provides an F1 score of 0.400 which is inferior to the performance under fixed generation.

### 3.3 Influence of Sequence Length on RNA Generation Quality

The quality of generated sequences is influenced by their length, as illustrated in Figure 2 (c). Longer sequence lengths expand the search space, requiring more time to generate optimal sequences. However, it is encouraging to note that shorter lengths yield strong results, particularly since therapeutic and targeted RNA sequence designs often necessitate shorter sequences [7].

## 4 Conclusion

In conclusion, our approach, RNA-DCGen, significantly enhances RNA sequence generation by incorporating dual constraints. This allows us to specify both the desired molecular properties and preserve critical segments of the sequence that hold biological significance. As a result, we can either generate complete sequences or focus on maintaining certain portions, ensuring that our outputs are structurally sound and functionally relevant.

A key advantage of RNA-DCGen lies in its flexibility. This method can be applied to any functional or structural conditional generation, provided that, we have the appropriate forward prediction dataset. Unlike other RNA generative tasks previously proposed that impose strict conditions on a particular property, RNA-DCGen can be adapted to accommodate various tasks as desired by the designer. To utilize this approach effectively, we require a dataset that maps sequences to specific properties, particularly at the nucleotide level. Additionally, fine-tuning an RNA language model (LLM) with this dataset is essential. By backpropagating the gradients from the LLM, we can perform a guided search that improves our ability to generate sequences that closely align with our desired characteristics, even in scenarios with limited data.

Moreover, the principles underlying RNA-DCGen can be readily extended to the design of proteins and other molecular structures. By applying the same dual-constraint framework, we can address similar challenges in protein engineering and molecular design. This adaptability positions RNA-DCGen as a valuable tool in bio-informatics, synthetic biology, and therapeutic design, paving the way for innovative solutions in sequence generation across diverse biological systems.

